# Microcompartment assembly around multicomponent fluid cargoes

**DOI:** 10.1101/2022.02.23.481520

**Authors:** Lev Tsidilkovski, Farzaneh Mohajerani, Michael F Hagan

**Affiliations:** Martin A. Fisher School of Physics, Brandeis University, Waltham, MA, 02453

## Abstract

This article describes dynamical simulations of the assembly of an icosahedral protein shell around a bicomponent fluid cargo. Our simulations are motivated by bacterial microcompartments, which are protein shells found in bacteria that assemble around a complex of enzymes and other components involved in certain metabolic processes. The simulations demonstrate that the relative interaction strengths among the different cargo species play a key role in determining the amount of each species that is encapsulated, their spatial organization, and the nature of the shell assembly pathways. However, the shell protein-shell protein and shell protein-cargo component interactions that help drive assembly and encapsulation also influence cargo composition within certain parameter regimes. These behaviors are governed by a combination of thermodynamic and kinetic effects. In addition to elucidating how natural microcompartments encapsulate multiple components involved within reaction cascades, these results have implications for efforts in synthetic biology to colocalize alternative sets of molecules within microcompartments to accelerate specific reactions. More broadly, the results suggest that coupling between self-assembly and multicomponent liquid-liquid phase separation may play a role in the organization of the cellular cytoplasm.

## I. INTRODUCTION

Compartmentalization is essential for many biological functions, including metabolism, cellular signaling, and genetic storage. While membrane-enveloped organelles are a prominent mode of compartmentalization in eukaryotic cells, it has become apparent that liquid-liquid phase separation^1–14^ and proteinaceous organelles^15–17^ play an important role in organizing the cytoplasm for all kingdoms of life. For example, in bacteria, certain metabolic pathways are enabled by bacterial microcompartments, which are organelles consisting of a large protein shell assembled around a dense complex of enzymes and reactants^18–25^. Other protein-shelled compartments include encapsulins^26,27^ and gas vesicles^26,28^ in bacteria and archea, and vault particles in eukaryotes^29^. From a practical perspective, there is great interest in exploiting principles and materials from biology to achieve similar nanoscale compositional control for synthetic biology and drug delivery applications. In particular, researchers have demonstrated the ability to target new molecules to microcompartment interiors and to transfect the systems into non-native organisms, including bacterial and plant cells, suggesting a basis for designing microcompartments as customizable nanoreactors (*e*.*g*.^22,30–43^).

Many of these cellular functions and biotechnology applications require compartments that assemble around multiple cargo species. For example, metabolism relies on achieving high local concentrations of the enzymes and reactants involved in a reaction cascade, while the formation of cellular signaling complexes requires colocalizing signaling proteins and their regulatory ligands^11,21,44^. Similarly, applications in synthetic biology require targeting multiple species to micro-compartment interiors (*e*.*g*.^22,24–26,30,31,34–43,45–52^). In particular, microcompartments could be exploited to increase the efficiency of arbitrary multi-enzyme cascades, if the species are encapsulated with controlled stoichiometry. Thus, understanding the factors that control the amount and composition of packaged cargo is essential for understanding both natural and re-engineered functions of microcompartments.

Previous experimental and modeling studies of micro-compartments and other shells provide an important starting point for understanding multicomponent encapsulation. The outer shells of microcompartments are roughly icosahedral, with diameters ranging from 40-400 nm, and assemble from pentameric, hexameric, and pseudo-hexameric (trimer-of-dimer) protein oligomers^18,19^. Recent atomic-resolution structures of small, empty microcompartment shells, as well as computational modeling^60–67^, have elucidated mechanisms that control the structure and size of micro-compartment shells. More broadly, previous modeling studies have shown the assembly of empty icosahedral shells^68–94^ and templating effects of a cargo consisting of a nanoparticle or RNA molecule on shell assembly^75,81–^. However, alternative models are needed for microcompartment encapsulation of multicomponent cargo complexes that do not have a specific size, structure, or composition.

Although previous experiments successfully targeted particular cargo species to microcompartment interiors^22,30^, the factors that control cargo coalescence and encapsulation remain incompletely understood. In some microcompartments, shell-cargo attractions are mediated by ‘scaffold proteins’ such as the CcmN protein in *β* -carboxysomes^120^, while in other systems core enzymes have a short ‘encapsulation peptide’ sequence that binds to the shell inner surface^26,31,^. Similarly, cargo-cargo attractions may be mediated by scaffolds, such as CcmM in *β* -carboxysomes^120^ or potentially through direct pair interactions between cargo molecules^36,37^.

Previous experimental and modeling studies suggest that the packaged cargo of microcompartments undergoes phase separation, either prior to or during assembly of the outer protein shell^60–62,6^. Furthermore, some microcompartments fulfill similar functions as liquid-liquid phase separated domains in cells — for example, the composition, structure, and function of the RuBisCO complex within carboxysomes has strong similarities to the pyrenoid, which is a liquid domain consisting of RuBisCO and other components that enables carbon fixation within plant cells^127–130^. It is therefore likely that some factors that control the composition of liquid-liquid phase separated domains in cells (*e*.*g*.^2,6,8,10,12–14,131–145^) also affect the composition of micro-compartment cargoes. However, in the case of microcompartment assembly, thermodynamic and kinetic factors resulting from coupling between liquid-liquid phase separation and assembly of the crystalline shell could have significant effects. Previous computational modeling of microcompartment assembly has focused on encapsulation of a single cargo species, driven either by direct cargo-cargo pair attractions^60,61,65^ or mediated by scaffolds^62^.

In this work, we build on these previous studies, by studying a minimal computational model for a microcompartment shell that assembles around a cargo containing two species. We investigate the amount and composition of the encapsulated cargo as a function of the affinities among these cargo species, and of the shell-cargo and shell-shell affinities. The simulation results suggest that the relative cargo-cargo affinities are the most important factor in determining the amount and composition of packaged cargo. However, within certain parameter ranges, varying the shell-cargo and shell-shell affinities provides an alternative means to sensitively control the packaged cargo. These effects are governed by a combination of thermodynamic and kinetic factors arising due to coupling between cargo coalescence and shell assembly.

## II. RESULTS AND DISCUSSION

### A. Model overview

We simulate the dynamics of microcompartment assembly in the presence of two cargo species, which we denote as ‘R’ and ‘G’. There are numerous control parameters in the microcompartment system, many of which have been explored in previous works^60–67^. Here, our objective is to focus on the effect of interaction parameters on cargo encapsulation. To avoid complexities associated with assembly into different shell geometries^61,62,65,66^, we use a minimal model for the shell developed in Perlmutter *et al*.^60^, for which the energy minimum corresponds to a *T* =3 icosahedral shell in the Caspar-Klug nomenclature^146^, comprising 12 pentamers and 20 hexamers (Section IV A and Fig. 1). Although our model is motivated by bacterial microcompartments, it is sufficiently general to describe other proteinaceous shells, including viruses and microcompartments that are reengineered to encapsulate designer cargoes (*e*.*g*.^22,30–39,41,46–48,50–52,147–156^).

**FIG. 1.**
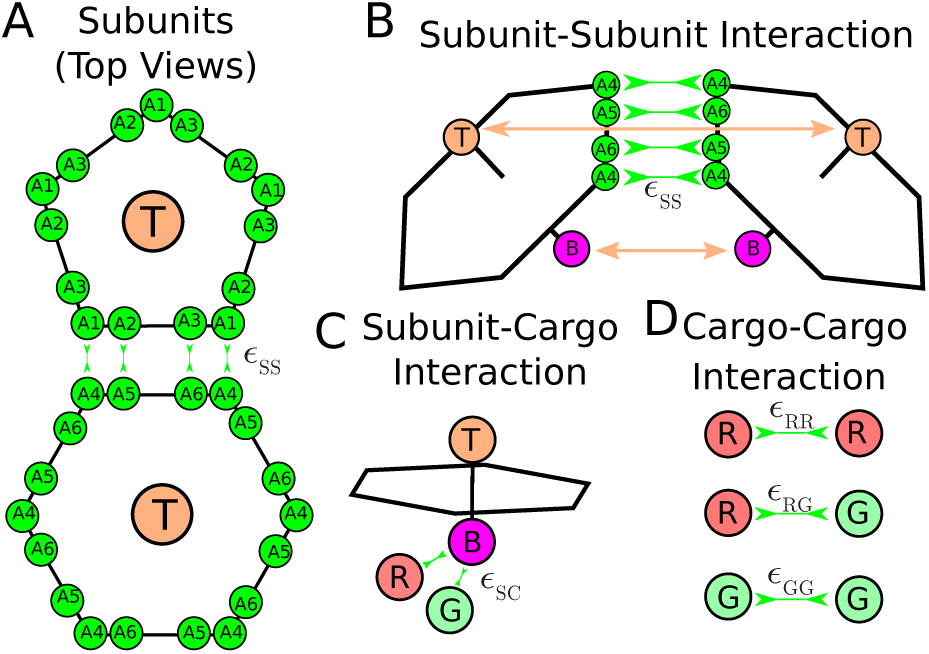
Description of the model. **(A)** Each shell subunit contains ‘Attractors’ (green circles) on the perimeter, that define the shape of the subunit. **(B)** Attractive interactions between Attractors drive subunit dimerization. Complementary pairs of Attractors are indicated by green arrows in (A) for the pentamer-hexamer interface and in (B) for the hexamer-hexamer interface. A combination of Top-Top (T) and Bottom-Bottom (B) repulsions controls the subunit-subunit angle in a complete shell. **(C)** Bottom pseudoatoms ‘B’ bind cargo molecules (shown as R and G). **(D)** The cargo molecules have attractive interactions between each other that depend on pair type.

#### Interaction parameter

Efficient assembly and cargo loading generically require three classes of interactions: interactions between shell subunits that drive shell assembly, interactions between shell subunits and one or more cargo species, and interactions between cargo particles that drive cargo coalescence^60^. While in some systems one or more of these interactions can be mediated by auxiliary proteins (*e*.*g*. the CCmM and CCmN proteins in *β* -carboxysomes), for simplicity we model all interactions as direct pair interactions in this work.

We have explored assembly dynamics over a range of the simulation control parameters that most strongly affect the amount and composition of encapsulated cargo — these are the interaction potential well-depths that control the binding free energy (affinity) between each type of cargo pair, *ε*_GG_, *ε*_RR_, and *ε*_RG_, as well as between shell-cargo and shell-shell pairs *ε*_SC_ and *ε*_SS_ (see Appendix B). To minimize the number of parameters, we assume that both cargo species have the same binding affinity to shell subunits; i.e., *ε*_SC_ is equal for R and G particles. For convenience, throughout this article we refer to the well-depths as ‘affinities’. We approximately relate *ε*_SC_ and *ε*_SS_ to the actual affinities in Appendix A and Table I. Throughout this article, all energy values are given in units of the thermal energy, *k*_B_*T*, and all lengths are given in units of the cargo diameter, *r*\*, which translates to *r*\*≈ 13 nm for carboxysomes (see section IV B).

**TABLE I.** The shell-shell binding affinity for the default affinity parameter values. The binding free energy and dissociation constant for hexamer dimerization (*g*_hh_, from Eq. (A1)), and from the shell-shell and shell-cargo interactions in a complete shell complex with *ε*_SC_ = 6 *k*_B_*T* (from Eq. (A3)). The standard state concentration is 1 M. _*_Does not include cargo-cargo contributions, which will increase (decrease) the effective shell dissociation constant 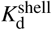 below (above) the threshold for cargo vapor-liquid coexistence (see Perlmutter et al.^60^)

|  |  |  |  |
| --- | --- | --- | --- |
| shell-shell affinity parameter | $\epsilon_{SS}/k_B T$ | 2.0 | 3.5 |
| hex-hex affinity | $g_{hh}/k_B T$ | 4 | −0.5 |
| hex-hex dissociation constant | $K_d^{hh}/\mu M$ | $7 \times 10^7$ | $6 \times 10^5$ |
| free energy per subunit in shell* | $\Delta g_{shell}/k_B T$ | −15 | −27 |
| dissociation constant from shell* | $K_d^{shell}/\mu M$ | 0.4 | $1 \times 10^{-6}$ |

We consider ranges of affinity parameters relevant to biological microcompartments (see section IV B for details). Except where mentioned otherwise, we consider two shell-shell affinities (*ε*_SS_*/k*_B_*T* = 2, 3.5), which are respectively below and above the transition between one-step and two-step assembly pathways (discussed in section II C). Both cases are below the threshold for assembly of empty shells (containing no cargo). Importantly, dimerization is very unfavorable, but the assembled shells are stable (see Table. I). The unfavorable dimerization affinities reflect the essential role that cargo plays in assembly at these shell-shell affinities, the cooperative nature of assembly, and the fact that a significant nucleation barrier is essential to avoid kinetic traps^60,80,89,157^. We primarily focus on a representative shell-cargo affinity *ε*_SC_ = 6 *k*_B_*T*, which corresponds to a free energy per shell subunit of *g*_SC_ ≈ − 14 *k*_B_*T*. Our simulations span the range of cargo-cargo affinities in the vicinity of vapor-liquid coexistence (*ε*vl ≈ 1.3 *k*_B_*T* at our simulated cargo concentration), but remain below the threshold for cargo crystallization at *ε*^fs^ ≈ 3 *k*_B_*T*. With *ε*^vl^ as a reference point, we denote cargo-cargo affinities as ‘weak’ for *ε*_RR_, : ≲ 1.1 *k*_B_*T*, ‘moderate’ for 1.2, : ≲ *ε*_RR_, : ≲ 1.4, and ‘strong’ for *ε*_RR_ ≳ 1.5 *k*_B_*T* (with analogous definitions for *ε*_GG_ and *ε*_RG_).

Our shell subunit and cargo concentrations map to approximately 1 *μ*M and 10 *μ*M respectively, which are reasonable based on quantities found within bacterial cells.

#### Simulated systems

There are multiple thermodynamic and kinetic factors which could influence the amount and composition of encapsulated cargo. Thermodynamic effects include the interplay between cargo-cargo and shell-cargo affinities, and the finite size of the encapsulated cargo globule. Kinetic effects arise from the fact that once a shell closes, even with relatively weak shell-shell affinities typical of productive assembly, exchange of shell subunits or cargo particles does not occur on experimentally relevant timescales. Thus, an assembling shell may trap its contents far out of equilibrium. To distinguish these effects, we begin by describing large simulations of cargo in the absence of shell subunits to approximate a bulk system (Section II B). We then describe the dynamical assembly of shells and cargo particles (Sections II C and II D). We compare the results of these finite-time dynamical simulations against simulations with assembled but permeable shells, which allow the encapsulated cargo to equilibrate with the bulk.

### B. Cargo phase behavior without shell subunits

We begin by briefly summarizing the bulk phase behavior of the cargo in the absence of shell subunits, as a reference point from which to understand how shell assembly can change cargo coalescence. We focus on parameter regimes relevant to our shell assembly simulations; more comprehensive descriptions of the phase behavior of a binary Lennard-Jones system can be found in *e*.*g*.^158–160^. In particular, we consider equal stoichiometry between R and G molecules and cargo-cargo affinities (*ε*_*i j*_ *<* 1.8 for *i j* = RR, GG, RG) that maintain the system in vapor and/or liquid phases, but are below the crystallization threshold (the vapor-fluid and crystallization transitions for a single species occur at at *ε*^vl^ = 1.3 and *ε*^fs^ ≈ 3). Within these limits, the phase behavior can be classified as follows: homogeneous demixed (no phase coexistence), phase separation of one cargo species (a dense phase rich in one cargo species in coexistence with a dilute phase containing an excess of the other), phase separation and mixing of both cargo species (coexistence between dilute and dense phases, with both species homogeneously mixed within both phases), and phase separation with de-mixing (coexistence between three phases — a dilute phase, a dense phase rich in R, and a dense phase rich in G). We consider weak, moderate, and strong R-G affinities: *ε*_RG_ = 1.0, 1.3, 1.6. For each of these, Fig. 2 summarizes the bulk phase behaviors as a function of the R-R and G-G affinities *ε*_RR_ and *ε*_GG_. Specifically, Figs. 2(A-C) compare the driving force for phase separation for each of the two species by showing the fraction of R particles *f*_R_ in the high-density phase (hereafter referred to as ‘globule’). Figs. 2 (D-F) quantify the extent of nonrandom compositional mixing within the high-density phase, *f*_UL_. Here we define the mean fraction of unlike neighbors in the first solvation shell around each particle as: 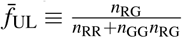, with *n*_*i j*_ the number of cargo neighbor pairs of species *i* and *j* with a neighbor pair defined as two particles with separations of *r <* 1.2. For random mixing, we expect *f*_rand_ = 2 *f*_R_(1 − *f*_R_), so the plots indicate the difference between randomness and the results: 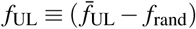.

**FIG. 2.**
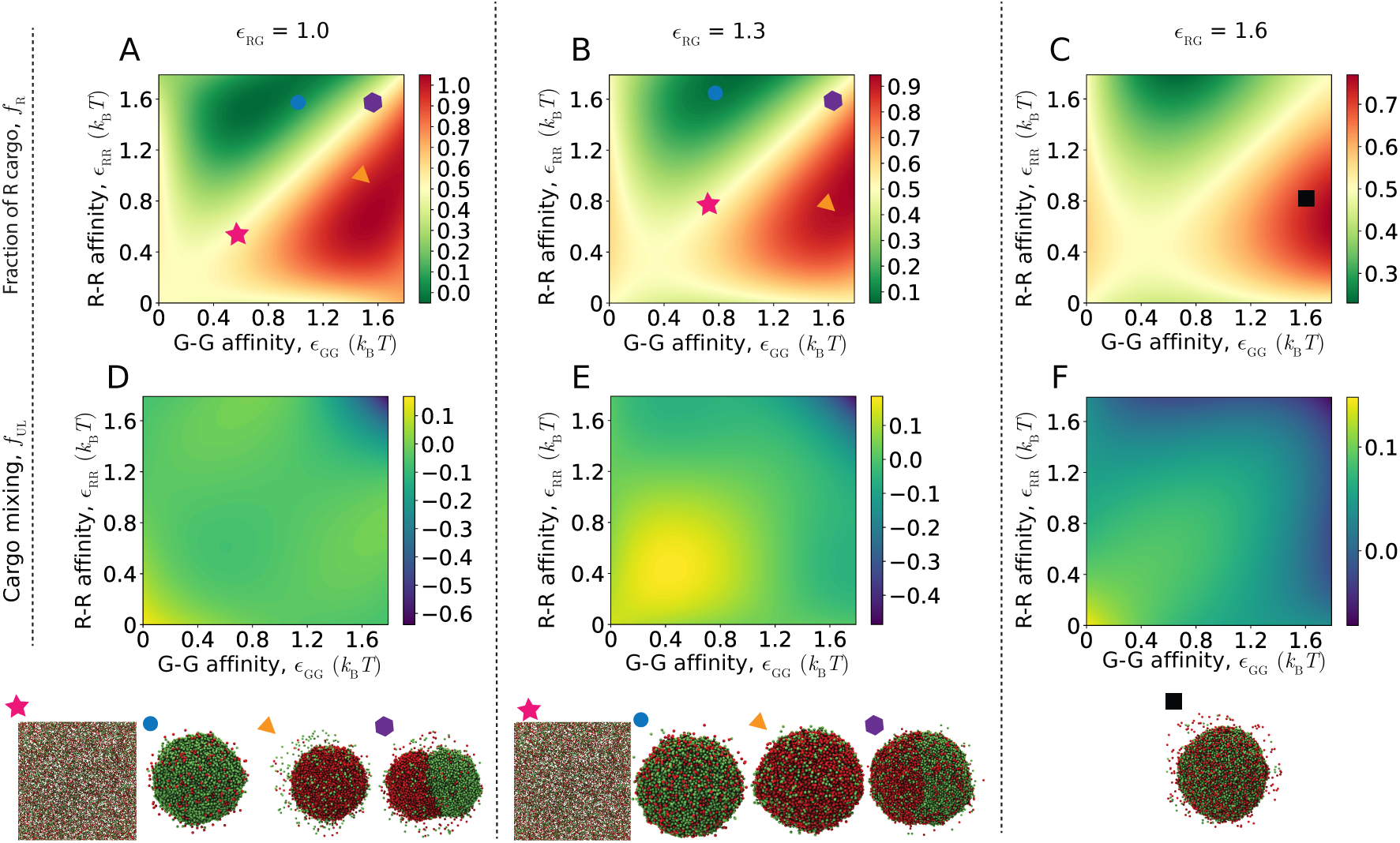
The bulk phase behavior of cargo particles (in the absence of shell subunits) as a function of their binding affinities. **(A-C)** The fraction of R particles in the high-density phase, *f*_R_, as a function of the R-R and G-G affinities for different R-G affinities: (A) weak *ε*_RG_ = 1.0, (B) moderate *ε*_RG_ = 1.3 and (C) strong *ε*_RG_ = 1.6. All energies are given in units of the thermal energy, *k*_B_*T*. The symbols on the plot identify corresponding snapshots below each plot that illustrate the phase behavior at the indicated value of *ε*_RG_. **(D-F)** For the same parameters, the fraction of unlike cargo particles in the first solvation shell relative to random mixing, *f*_UL_, defined in the text. Each simulation contains 46,938 total cargo particles, with equal composition of R and G.

For weak R-G affinities (*ε*_RG_ = 1.0, Fig. 2A) the system exhibits either no phase separation, coexistence between a phase rich in one species and a dilute phase, or coexistence among a dilute phase and separate phases for each species. In the latter case, the R-rich and G-rich phases are attracted to each other by the weak but non-zero R-G affinities, leading to formation of one globule with an interface separating the two phases. The interface between these two phases becomes more diffuse with decreasing *ε*_RR_ and *ε*_GG_, and/or increasing *ε*_RG_ because the surface tension between the two domains decreases. For moderate R-G affinities *ε*_RG_ = 1.3, Fig. 2B) we observe no phase separation, coexistence between a phase rich in one species and a dilute phase, or coexistence between a dilute phase and a dense phase containing both species.

For strong R-G affinities (*ε*_RG_ = 1.6, Fig. 2C) we always observe mixing of both species within the dense phase, with the fraction of R or G particles depending on the relative magnitudes of *ε*_RR_ and *ε*_GG_. Since the primary driving force for phase separation is *ε*_RG_ in this case, fluctuations in *f*_R_ and *f*_UL_ are smaller than for the other systems.

### C. Assembly pathways

We now consider how cargo and shell affinities affect the dynamics of assembly and cargo encapsulation. In previous work, simulations with a single cargo species showed that assembly pathways are strongly affected by the net strength of cargo-cargo affinities^60^. In particular, at our simulated volume fraction of 0.003, weak cargo-cargo affinities (*ε*_RR_ *<* 1.7) and/or the higher shell-shell affinities (*ε*_SS_ *>* 2.5) lead to one-step pathways in which shell assembly is concomitant with cargo coalescence, while strong cargo-cargo and lower shell-shell affinities (*ε*_RR_ *>* 1.7 and *ε*_SS_ *<* 2.5) allow two-step pathways in which a cargo globule coalesces, followed by disordered adsorption of shell subunits onto the globule surface and then cooperative rearrangement into an assembled shell. Other interaction combinations, such as high cargo-cargo and shell-shell affinities lead to pathways in between these two extremes.

The case of two cargo species exhibits qualitatively similar classes of assembly pathways. Fig. 3A,B show snapshots from typical simulations at shell-shell affinity *ε*_SS_ = 3.5, leading to one-step assembly pathways. In Fig. 3A the cargo-cargo affinities are uniform and relatively strong *ε*_RR_ = *ε*_GG_ = *ε*_RG_ = 1.7, leading to rapid and simultaneous coalescence of cargo and shell assembly, resulting in a uniform mixture of both cargo types within the complete shell. In Fig. 3B the R-R affinities (*ε*_RR_ = 1.7) are stronger than the G-G and R-G affinities (*ε*_GG_ = *ε*_RG_ = 1.3), leading to simultaneous assembly and coalescence of a nearly pure R domain.

**FIG. 3.**
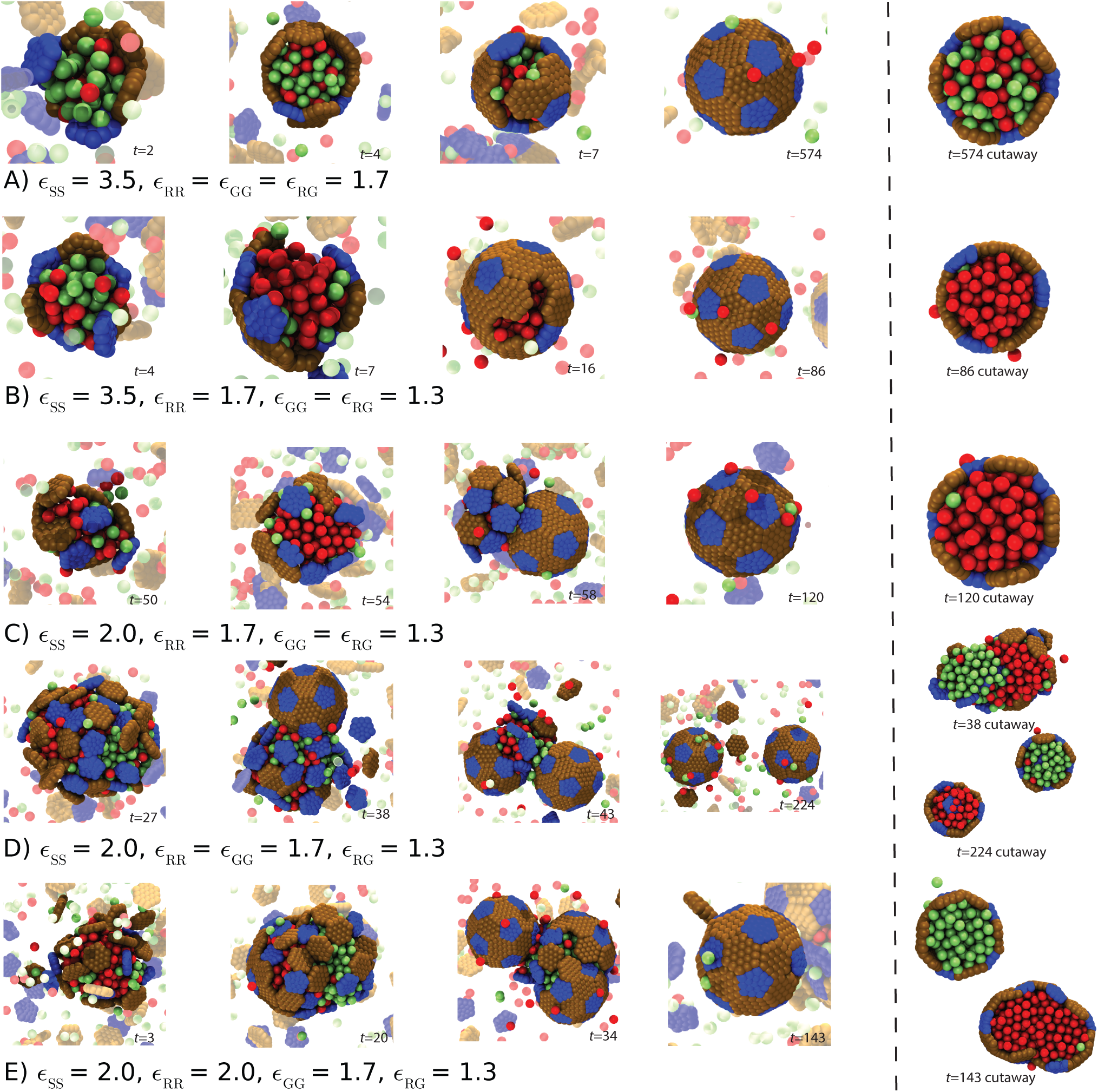
Assembly pathways. Snapshots from simulation trajectories illustrate the classes of assembly pathways discussed in the text. In each row, the first four snapshots show frames at indicated time points (in units of the nondimensional time *τ*), while the last snapshot is a cutaway view of the shell to show the encapsulated cargo. The parameters characterizing binding affinities are listed under each row (shell-shell, *ε*_SS_; shell-cargo, *ε*_SC_; R-R and G-G, *ε*_RR_ and *ε*_GG_; and R-G *ε*_RG_. **(A)** All affinities are relatively strong, driving rapid and simultaneous coalescence of both cargo types and shell assembly (one-step assembly). The encapsulated cargo is uniformly mixed. **(B)** At *ε*_SS_ = 3.5 *k*_B_*T*, high *ε*_RR_ but moderate *ε*_GG_ and *ε*_RG_ lead to one-step assembly and coalescence of R cargo, with G cargo essentially excluded from the shell. **(C)** At *ε*_SS_ = 2 *k*_B_*T*, strong *ε*_RR_ but moderate *ε*_GG_ and *ε*_RG_, lead to two-step assembly around an almost pure R domain. **(D)** Strong *ε*_GG_ and *ε*_RR_ and moderate *ε*_RG_ drive two-step assembly, with coupling between shell closure and cargo compositional fluctuations, leading to encapsulation of nearly pure R and G cargo domains in separate shells. The initial globule that becomes separated is shown in the top cutaway view, while the assembled shells are shown in the bottom view. **(E)** Very strong *ε*_RR_, strong *ε*_GG_, and moderate *ε*_RG_ lead to two-step assembly around an almost pure G domain. The strength of the *ε*_RR_ interaction prevents the subunits from successfully encapsulating the globule of R cargo, while the G cargo is able to bud off similarly to **(D)** and form a properly assembled shell. The cutaway shows the state of the assembled shell and G cargo domain within as well as the R cargo globule that is unable to properly close. Supplemental videos, corresponding to each of the respective trajectories are included in the SI (Multimedia view).

Fig. 3C-E show trajectories with a weaker shell-shell interaction of *ε*_SS_ = 2.0, for which shell nucleation is slow and we observe two-step assembly pathways. In Fig. 3C, the cargo-cargo affinities are strongest for R-R, and the assembly pathway closely resembles that for a single species — a domain of nearly pure R particles condenses, with subsequent assembly and encapsulation by the shell (Multimedia view).

We observe a more interesting coupling between cargo composition and shell assembly for strong R-R and G-G affinities (*ε*_RR_ = *ε*_GG_ = 1.7) and moderate R-G affinities *ε*_RG_ = 1.3 (Fig. 3D). The cargo initially coalesces into a globule that is mixed, albeit with significant compositional fluctuations, consistent with the bulk cargo phase behavior under these parameters (the 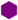 symbol in Fig. 2B). At these parameters, the domain is large enough to form two shells and shell closure couples to compositional fluctuations and occurs preferentially in the vicinity of an interface between R-rich and G-rich domains. In particular, we observe closure at the interface and consequently purified cargo encapsulation, meaning that the encapsulated cargo within an individual shell is nearly pure R or G.

The origins of coupling between shell closure and compositional fluctuations can be understood from the competing forces that govern shell closure. For the shell to close, curvature of the assembling shell must first generate the formation of a ‘neck’ of cargo, i.e. a narrow region of cargo particles that connects a shell assembly to the remainder of the globule or to other assemblies (*e*.*g*. see Fig. 3D at *t* = 43 and Fig. 3E at *t* = 34). Then, additional shell subunits must assemble to close the shell, causing the neck to break and the completed shell to separate from the remainder of the globule (Fig. 3D at *t* = 224 and Fig. 3E at *t* = 143). The processes of forming and then breaking the neck increase the total cargo interfacial area and correspondingly reduce the number of cargo-cargo interactions. Thus, they are accompanied by a free energy barrier with a height that increases with cargo surface tension, and correspondingly increases with cargo-cargo affinity. In Fig. 3E, the R-R affinities are stronger (*ε*_RR_ = 2.0), resulting in a larger barrier for closure within the R-rich portion of the globule, and thus a complete shell forms only around G particles.

#### Shell closure locks the encapsulated cargo composition out of equilibrium

After a shell closes, fluctuations of subunits large enough to allow cargo particles to escape are extremely rare because all subunits have their maximum number of interactions. Thus, the cargo composition is essentially fixed once the shell completes. However, as noted in Section II D the cargo remains fluid in the range of cargo-cargo affinities that we focus on, and thus the cargo can spatially reorganize inside of the shell even after its completion.

### D. Factors that control the composition of encapsulated cargo

#### 1. Cargo-cargo affinity

Under most conditions the cargo-cargo affinities have the strongest effect on the amount and composition of encapsulated cargo. We begin by considering one-step assembly pathways obtained with moderate shell-cargo affinities *ε*_SC_ = 6.0 and the higher shell-shell affinities *ε*_SS_ = 3.5. These conditions allow assembly over a broad range of cargo-cargo affinities.

Fig. 4 shows observables that characterize cargo properties, as well as the assembly yield, quantified as the fraction of subunits within complete shells (*f*_s_, Fig. 4D). Because the most interesting variations occur at moderate R-G affinities, we present most results in this figure with varying R-R and G-G affinities at fixed *ε*_RG_ = 1.3. First, Fig. 4C shows that the amount of packaged cargo increases monotonically with the net encapsulation driving force; *i*.*e*., increasing either or both of *ε*_RR_ and *ε*_GG_ raises the number of encapsulated cargo particles *N*_C_, which asymptotically approaches the value corresponding to crystalline density (*N*_C_ ≈ 150)^60^. This demonstrates that cargo-cargo cohesive interactions are essential to obtain full shells^60^. Importantly though, *N*_C_ does not reach crystalline density until cargo-cargo affinities nearly reach the bulk crystallization threshold *ε*^fs^ ≈ 3 *k*_B_*T*. Ref. ^60^ found that well-formed shells cannot assemble when the cargo crystallizes, because the solid cargo cannot be deformed by assembly subunits, thus preventing shell closure. They also measured radial distribution functions for a single cargo species system, which showed that confinement by the shell imposes layering on the encapsulated cargo due to steric effects, but that the cargo remains fluid. The same effect is present for the bicomponent system studied in the present work, since the two cargo species differ only in their cargo-cargo affinities.

**FIG. 4.**
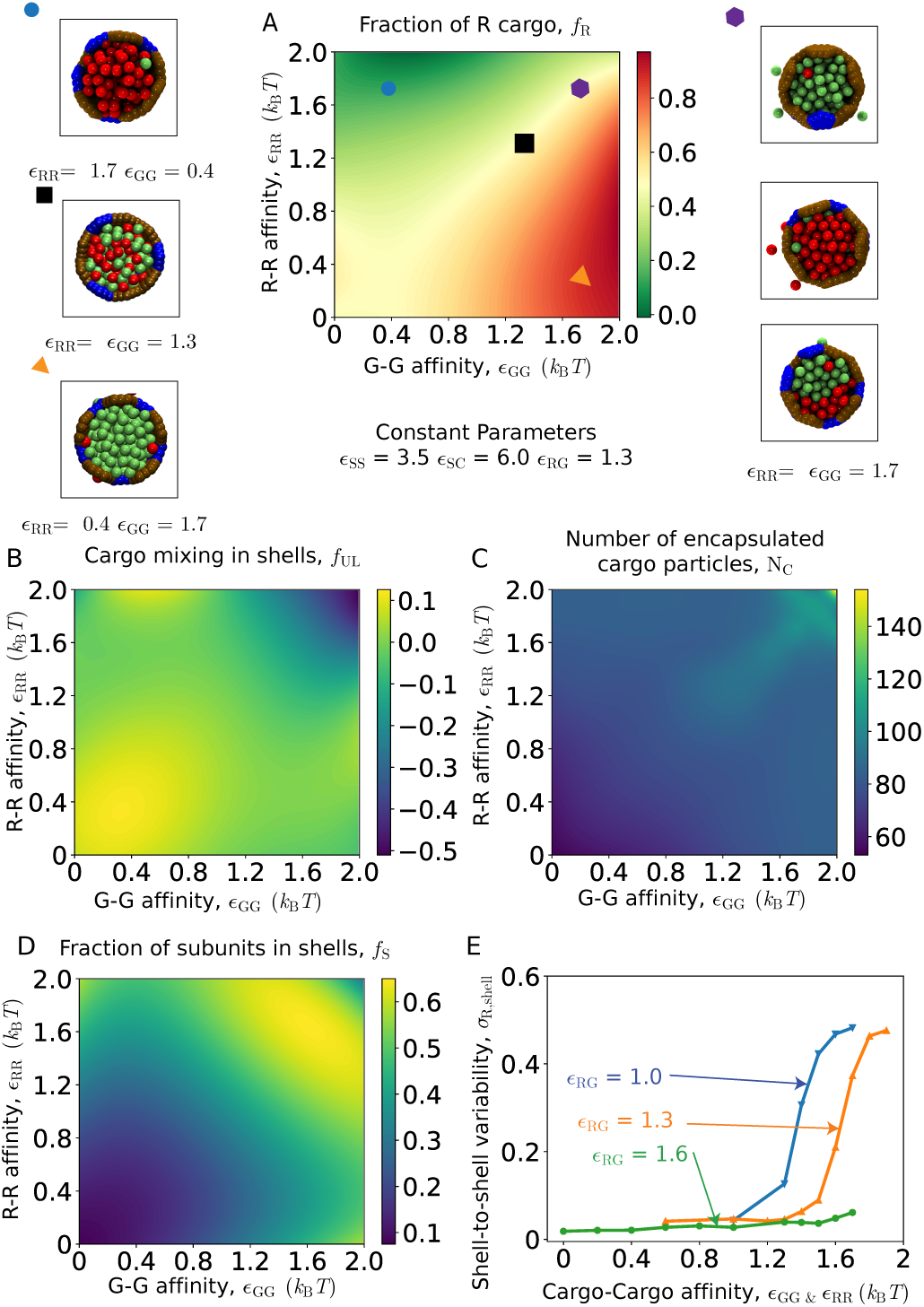
Effect of cargo-cargo affinities on encapsulation for one-step assembly pathways. **(A-C)** Characterization of the encapsulated cargo as a function of the G-G and R-R affinities, for fixed R-G affinity *ε*_RG_ = 1.3. **(A)** The fraction of R particles averaged over all shells *f*_R_ to characterize the average composition of encapsulated cargo. Snapshots show cutaway views of assembled shells, to illustrate the cargo morphology at corresponding symbols on the plot. **(B)** The fraction of unlike cargo particles in first solvation shells, *f*_UL_ averaged over all shells. **(C)** The mean number of encapsulated cargo particles per shell, to indicate cargo loading efficiency. **(D)** The fraction of subunits in complete shells, revealing the effect of cargo-cargo affinities on shell assembly. **(E)** The shell-to-shell variability of cargo composition, *σ*_R,shell_, defined as the standard deviation of *f*_R_ over shells, is shown as a function of the R-R and G-G affinities for three indicated values of *ε*_RG_. High values of *σ*_R,shell ≳_ 0.4 indicate that different shells have encapsulated pure domains of respectively R or G cargo species. To simplify the presentation, we only consider equal R-R and G-G affinities, *ε*_RR_ = *ε*_GG_. For (A)-(E) other parameter values are *ε*_RG_ = 1.3, shell-cargo affinity *ε*_SC_ = 6.0, and shell-shell affinity *ε*_SS_ = 3.5.

Next, to determine the relative loading of the two cargo species, Fig. 4A shows the fraction of R species averaged over all shells, *f*_R_, as a function of *ε*_GG_ and *ε*_RR_. Further, to characterize the arrangement of cargo particles within shells, Fig. 4B shows the fraction of unlike particles in the first solvation shell (*f*_UL_). The snapshots surrounding Fig. 4A illustrate typical arrangements of the encapsulated cargo in each regime. Comparing Figs. 4A,B with results from the bulk cargo system in the absence of microcompartment shells (Figs. 2B,E), shows that the composition and mixing of the two cargo species within shells is similar to the case of bulk cargo coalescence for this parameter regime. However, we will see below that shell assembly has a more significant effect on cargo properties in other parameter regimes.

Fig. 5 compares these results to two-step assembly pathways that occur for lower shell-shell affinities, *ε*_SS_ = 2.0. Since we observe significant assembly only for a narrow range of strong cargo-cargo affinities (Fig. 8, discussed next), we present these results as a function of *ε*_RR_ for fixed *ε*_GG_ = 1.7. We see that average cargo loading and composition are qualitatively similar between the two cases, although there are quantitative differences revealing deviations from equilibrium, which we consider in more detail in Section II D 2.

**FIG. 5.**
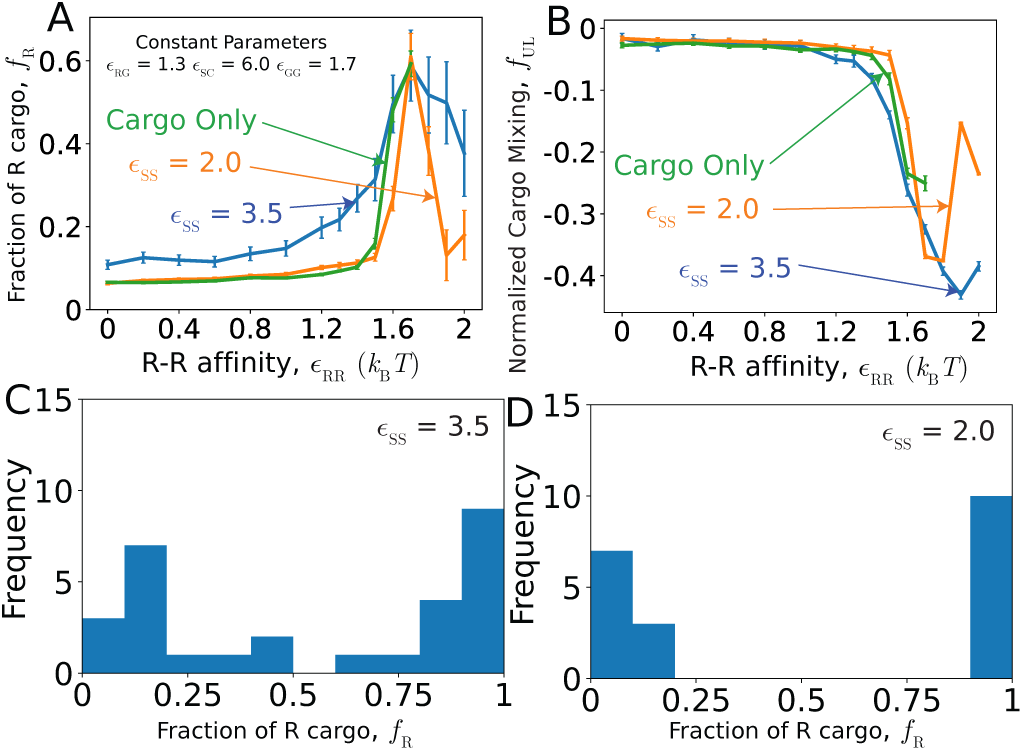
Comparison of cargo encapsulation for one-step and two-step pathways. The fraction of R cargo in completed shells *f*_R_ **(A)** and the fraction of unlike cargo particles in first solvation shells *f*_UL_ **(B)** are shown as a function of *ε*_RR_ for the lower and higher shell-shell affinities *ε*_SS_ = 2.0, and *ε*_SS_ = 3.5. Other parameters are *ε*_GG_ = 1.7, *ε*_RG_ = 1.3, and *ε*_SC_ = 6.0. **(C, D)** show histograms of cargo composition within individual shells to illustrate shell-to-shell variability, for the peak in panel (A) at *ε*_RR_ = *ε*_GG_ = 1.7 and *ε*_RG_ = 1.7, for (C) one-step and (D) two-step pathways.

**FIG. 6.**
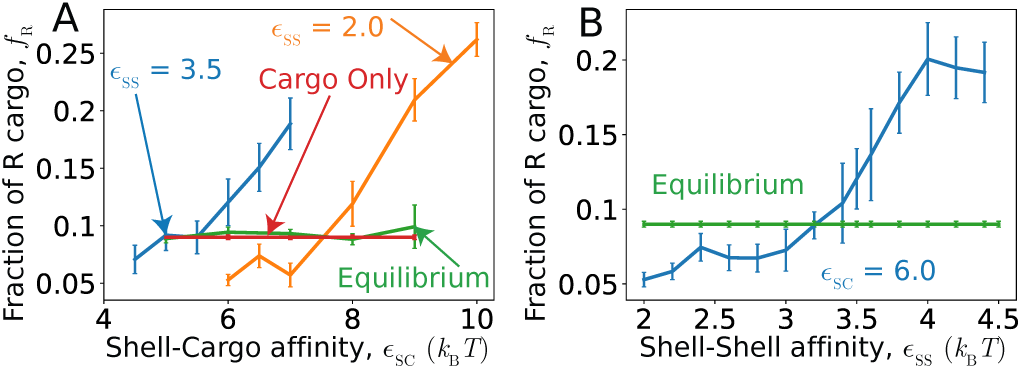
Effects of shell-cargo and shell-shell affinity on cargo composition reveal thermodynamic and kinetic influences on encapsulation. **(A)** The fraction of encapsulated R particles *f*_R_ as a function of shell-cargo affinity *ε*_SC_, for shell-shell affinities *ε*_SS_ = 2.0, and *ε*_SS_ = 3.5. To assess the importance of dynamics on these results, *f*_R_ is also shown for the ‘equilibrium’ simulations in which cargo can exchange between shell interiors and the bulk. Other parameters are *ε*_GG_ = 1.7 and *ε*_RR_ = *ε*_RG_ = 1.3. **(B)** Cargo composition *f*_R_ as a function of shell-shell affinity *ε*_SS_, at a constant shell-cargo affinity of *ε*_SC_ = 6.0. Other parameters are *ε*_RR_ = 1.3, with *ε*_GG_ = 1.7 and *ε*_RG_ = 1.3. The ‘equilibrium’ results are also shown, as a straight line since they do not depend on *ε*_SS_.

However, the composition of cargo within individual shells can differ significantly from the mean. Fig. 4E shows the shell-to-shell variability of cargo composition, *σ*_R,shell_, defined as the standard deviation of the fraction of R particles within each shell: 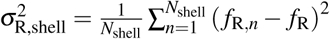. Because compositional fluctuations depend strongly on the ratio of *ε*_RR_ and *ε*_GG_ to *ε*_RG_, we present the results as a function of equal R-R and G-G interactions (*ε*_RR_ = *ε*_GG_), for weak, moderate, and strong *ε*_RG_. In each case, as the R-R and G-G affinities exceed *ε*_RG_ there is a transition from shells containing uniformly mixed cargo to nearly pure domains of R or G particles respectively within different shells, corresponding to large *σ*_R,shell_.

Interestingly, this variability arises in different ways depending on the class of assembly pathways. To illustrate this difference, Fig. 5C,D show histograms of cargo compositions measured within individual shells for one-step and two-step pathways respectively. These parameters correspond to the maximum in Fig. 5A, where *f*_R_ and *f*_UL_ are approximately equal for the two pathways, but the mechanisms underlying the cargo separation differ. In one-step pathways, as shown in Fig. 3B, a compositional fluctuation in the initial cargo globule nucleus leads to preferential coalescence of the same cargo type as shell assembly proceeds, with the shell eventually closing around a nearly pure domain. However, for these parameters we do occasionally (∼ 5%) observe shells that encapsulate two domains separated by an interface (see Fig. 5C). In contrast, two-step assembly pathways form purified shells via the compositional fluctuation mechanism discussed in Section II C and shown in Fig. 3D. This leads to highly purified cargo globules within individual shells (see Fig. 5D).

##### Cargo-cargo affinities also affect shell assembly

While our focus in this article is on factors that control cargo encapsulation, it is important to note that the driving force for cargo cohesion in turn affects the extent and robustness of shell assembly. Figs. 4D and Fig. 8A show the fraction of subunits in complete shells *f*_s_ as a function of cargo-cargo affinities for *ε*_SS_ = 2.0 and *ε*_SS_ = 3.5 respectively. We observe that assembly occurs over a broad range of cargo-cargo affinity values for the higher shell-shell affinity (*ε*_SS_ = 3.5, Fig. 4D), but for the lower shell-shell affinity (*ε*_SS_ = 2.0, Fig. 8) significant assembly requires at least one strong cargo-cargo affinity; *i*.*e*., *ε*_*i j*_ ≥ 1.7 for at least one of *i j* ∈ {RR, GG, RG }. This result indicates that nucleation of shell assembly for *ε*_SS_, : ≲ 2.5 requires adsorption onto a cargo globule and thus requires that cargo-cargo affinities are sufficient to drive cargo phase separation. This observation is consistent with results from simulations with a single cargo species^60^.

#### 2. Shell-shell and shell-cargo affinities affect cargo packaging through equilibrium and non-equilibrium mechanisms

Although the cargo-cargo affinities most strongly affect the amount and properties of encapsulated cargo, the shell protein affinities (*ε*_SC_ and *ε*_SS_) can also modulate cargo encapsulation. These effects depend on an interplay between thermodynamic and kinetic effects. The assembly trajectories in our simulations begin from an initial condition of dispersed subunits and cargo that are significantly out of equilibrium. Moreover, once a shell closes, the timescale for reconfiguration to another shell morphology or exchange of the encapsulated cargo is much longer than assembly timescales since all subunits have their maximum number of interactions. Therefore, even though the dynamics satisfy microscopic reversibility, trajectories are not guaranteed to approach equilibrium on the finite timescales of simulations and experiments^157^.

To distinguish between equilibrium and kinetic effects, we compare the results of our assembly simulations to the alternative ‘equilibrium’ or permeable-shell system described above, in which shells are completely assembled but are made permeable to cargo particles, thus allowing the encapsulated cargo to fully equilibrate with cargo particles in solution (see section IV). The amount and composition of encapsulated cargo in these simulations is thus independent of assembly kinetics (and shell-shell affinities). Fig. 7 compares the results of assembly trajectories at parameters leading to one-step assembly (corresponding to Fig. 4) with the equilibrium simulations. Over much of parameter space the amount and compositions of encapsulated cargo are similar between the two systems, indicating that trajectories are near equilibrium during assembly. This can be understood since the diffusional exchange of cargo particles between the partially encapsulated cargo globule and bulk is rapid compared to shell growth under those conditions.

**FIG. 7.**
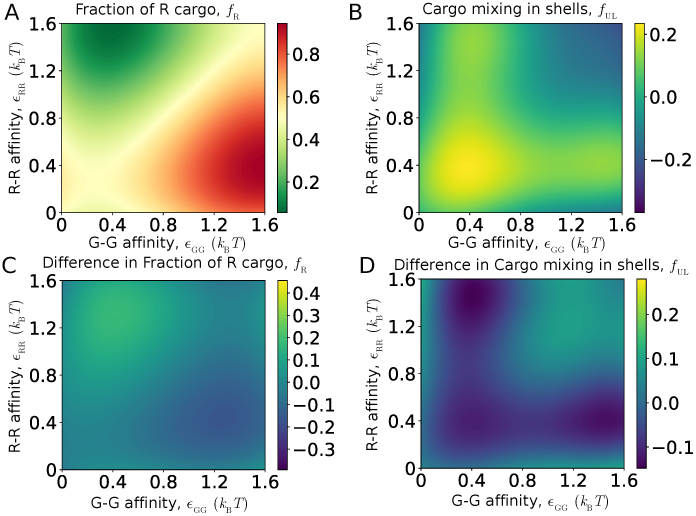
Comparison of encapsulated cargo between equilibrium and dynamical simulations. **(A**,**B)** The fraction of R particles **(A)** and mixing **(B)** are shown as a function of R-R and G-G affinities. Other parameters are *ε*_RG_ = 1.3 and *ε*_SC_ = 6.0, as in Fig. 4. **(C**,**D)** Difference between cargo encapsulation in dynamical assembly simulations and equilibrium. The plots show the differences between the quantities *f*_R_ and *f*_UL_ for the dynamical simulations Fig. 4 (with *ε*_SS_ = 3.5) and equilibrium simulations (panels (A,B) of this figure).

**FIG. 8.**
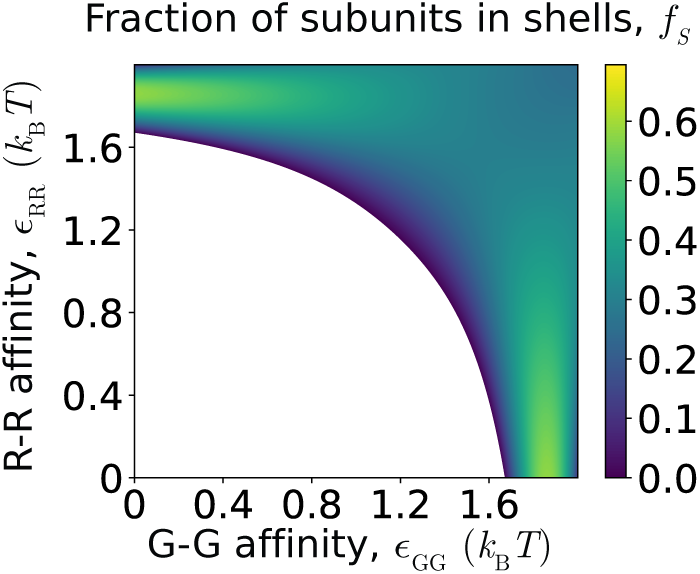
Fraction of shell subunits in complete shells in the two-step assembly pathway regime, *ε*_SS_ = 2.0 and *ε*_RG_ = 1.3.

However, we do observe significant deviations from equilibrium cargo encapsulation under certain parameter ranges. These effects are strongly dependent on the shell-shell and shell-cargo affinities, since they are key determinants in the extent to which assembly is out of equilibrium.

For example, Fig. 5A shows the dependence of *f*_R_ on *ε*_RR_ for one-step (*ε*_SS_ = 3.5) and two-step (*ε*_SS_ = 2.0) assembly pathways. For moderate cargo-cargo affinities *ε*_RR_, : ≲ 1.5, the two-step assembly and equilibrium results closely agree, while the one-step simulations result in higher *f*_R_. This difference arises because the initial globule is enriched in R particles (since globule nucleation is driven in part by shell-cargo attractive interactions that are agnostic to particle type), and the rapid one-step assembly pathways do not allow the cargo globule time to fully equilibrate with the bulk. In contrast, the slow onset of nucleation and assembly in the two-step pathways allows globule equilibration. Interestingly, the situation reverses at high *ε*_RR_ ≳ 1.6, where there is an abrupt drop in *f*_R_ for the two-step case. This arises because the strong R-R affinities favor segregation of the two cargo species and a high surface tension that inhibits closure of a shell around R particles. A comparable trajectory is shown in Fig. 3E. Notice that a shell successfully assembles around the cargo particles with weaker affinities (G), but only a malformed structure forms around the R globule because the shell subunits are unable to close around a commensurate-sized piece of the globule. Note that we do not show the permeable-shell results for such high *ε*_RR_ because the strong cargo-cargo affinities result in a cargo globule that extends outside of the shell, which is an artifact of the permeable-shell condition imposed in those simulations.

We also observe that the extent of cargo mixing within shells (Fig. 5B) decreases with increasing *ε*_RR_. This is partially an equilibrium effect due to the increasing drive for cargo segregation, especially as the R-R and G-G affinities exceed the R-G affinity (*ε*_RG_ = 1.3). However, while the twostep results are close to equilibrium, mixing is further reduced for the one-step assembly trajectory. This effect can be attributed to the coupling between composition fluctuations and interfacial effects within the globule during the initial stage of the two-step pathway described in section II C and Fig. 3D. Due to interfacial tension, the boundary between the two domains provides a ‘weak point’ for insertion of shell subunits, and thus shell closure tends to occur in the vicinity of the interface. The closing shell becomes enriched in the cargo species that dominates the domain on the same side of the interface as the assembling shell; this composition is then effectively locked in by shell closure.

Having compared the differences in cargo encapsulation between the two classes of assembly pathways, we now consider parameter values between these two extremes. Figs. 6A and 6B respectively show the effects of shell-cargo and shell-shell affinities on the composition of encapsulated cargo. Here, we have focused on the case for which cohesive interactions between pairs of G particles are stronger than for R-R and R-G, *ε*_GG_ = 1.7 and *ε*_RR_ = *ε*_RG_ = 1.3, for which the equilibrium and bulk composition of R particles is about 20%. Fig. 6A shows that for weak shell-cargo affinities, the encapsulated fraction of R particles is significantly smaller than the equilibrium value, but that it monotonically increases with *ε*_SC_. The trend is the same for both shell-shell affinities, except that higher shell-cargo affinities are required to observe assembly for lower shell-shell affinities. Similarly, Fig. 6B shows that *f*_R_ increases monotonically with shell-shell affinity. These trends can be understood as follows. At low *ε*_SC_, the cargo-cargo affinities are the primary driving force for encapsulation, which at these parameters favors encapsulation of G cargo and thus small *f*_R_. The fact that *f*_R_ in the dynamical assembly simulations is smaller than that observed in the bulk cargo and equilibrium simulations suggests that the initial globule that nucleates is enriched in G, and that the shell closes before there is sufficient time for the globule composition to equilibrate with the bulk. However, as *ε*_SC_ or *ε*_SS_ increases, the strong shell-cargo affinities enhance nucleation, allowing a cargo globule to form with a composition closer to the bulk composition, *f*_R_ = 0.5.

## III. CONCLUSIONS

We have performed equilibrium and dynamical simulations to investigate the encapsulation of multicomponent fluid cargoes by self-assembling microcompartments. To elucidate mechanisms that control the amount and composition of cargo encapsulation, we compared these results against simulations of cargo coalescence in the absence of microcompartment assembly.

Our simulation results show that the binding affinities among the different cargo species are the strongest determinant of the assembly pathway, as well as the amount, composition, and spatial organization of cargo within assembled shells. Analogous to the case of encapsulation of a single component cargo^57,60,61,120,122,123,161^, the simulations exhibit two broad classes of assembly pathways. Relatively strong cargo-cargo affinities and low shell-shell affinities lead to twostep assembly, in which the cargo first forms a globule, followed by adsorption and assembly of the shell around the globule. Weaker cargo-cargo affinities and higher shell-shell affinities drive one-step assembly, in which cargo coalescence and shell assembly occur simultaneously. However, the bi-component cargo allows for more diverse assembly pathways and morphologies, depending on when and if the two components mix within the encapsulated globule. In particular, we observe assembly pathways ranging from coalescence and encapsulation of a nearly pure globule of a single cargo species, to coalescence of a phase-separated globule with each phase becoming encapsulated in different shells, to encapsulation of a globule consisting of a uniform mixture of both species. Correspondingly, the assembled shells can contain a variety of cargo compositions and morphologies, ranging from highly enriched in a single species to nearly uniform mixing of the two species. In regimes where both cargo species are encapsulated, the two species may be uniformly mixed within each shell, or separated, with each individual shell containing an encapsulated cargo that is nearly pure in one or the other species.

While the relative strengths of the different cargo-cargo binding affinities (*ε*_RR_, *ε*_GG_, and *ε*_RG_) determine cargo composition and mixing, the net cargo-cargo affinity most strongly influences the amount of cargo that is encapsulated. In particular, for low cargo-cargo affinities we observe poor cargo loading (*e*.*g*. shells that are nearly empty except for a layer of cargo at the surface). Further, depending on the strength of the subunit-subunit affinities, there is a threshold value of cargo-cargo affinities below which assembled microcompartments are either unstable or fail to nucleate on relevant timescales (Figs. 4D and 8).

While cargo composition and mixing within assembled shells is qualitatively similar to the behavior of cargo coalescence in bulk (in the absence of microcompartments) in many parameter regimes, our simulations also identify parameter regimes in which the shell-cargo and shell-shell binding affinities can be manipulated to control cargo encapsulation (*e*.*g*. Fig. 5 and Fig. 6). A combination of thermodynamic and kinetic mechanisms underlie these effects. In particular, we see a marked difference between the properties of the encapsulated cargo in shells that assemble by one-step or two-step pathways. When cargo-cargo affinities are asymmetric, leading to preferential encapsulation of one species, weaker shell-cargo and shell-shell affinities and two-step assembly pathways tend to enhance the degree of preferential encapsulation leading to more purified cargo. This behavior reflects the fact that the cargo globule that nucleates in the first step tends to be enriched in the species with higher affinities, and, if assembly occurs more rapidly than the process of cargo exchange with bulk, shell closure locks in this out-of-equilibrium composition. Similarly, when the set of cargo-cargo affinities places the system close to ternary phase coexistence (a dilute phase and two dense phases respectively rich in either cargo species), two-step assembly pathways can enhance separation of the cargo species. This result can be explained at least in part by the tendency of assembling shells to close in the vicinity of transient interfaces between domains of the two cargo species (Fig. 2).

The latter predictions can be tested in microcompartment systems by performing mutagenesis of shell protein-protein binding interfaces (to alter shell-shell affinities) or ‘encapsulation peptides’ that mediate shell-cargo interactions^162–165^. Similarly, cargo-cargo affinities may be tuned through mutagenesis, but they are less well understood at present. Depending on the system, cargo-cargo affinities may arise from direct pair interactions, scaffold-mediated interactions, or a combination of the two^31,36,39,121,122,126,166,167^.

*Outlook*. It will be interesting to extend our model to explicitly incorporate scaffold-mediated cargo-cargo and shell-cargo interactions, for example as Mohajerani *et al*.^62^ did for a single-component cargo. More broadly, the predictions from our models can be extended beyond microcompartments, for example to re-engineered viral capsids^46–48,51,147–155^ or synthetic capsids constructed via protein engineering (*e*.*g*.^168–170^) or DNA origami^171^ to assemble around specific cargoes. In some of these cases, the shell-cargo interactions are driven by ‘scaffolds’ attached to the inner surface of the capsid. For example, many virus capsid proteins have flexible peptide tails that drive RNA encapsulation^172–174^, while Sigl et al.^171^ designed DNA origami capsid subunits with single-stranded DNA oligomers that bound to complementary strands on cargo particles. Previous simulations of microcompartments^62^ and virus assembly^106,175–178^ suggest that the properties of such scaffolds can be designed to manipulate the spatial organization of the cargo. Finally, it will be important to allow for shell assembly into different geometries, to study coupling between shell size and cargo composition.

## IV. METHODS

### A. Computational model

We consider a minimal model for microcompartment assembly and encapsulation of two species of cargo particles, denoted as ‘R’ and ‘G’. Microcompartment shells assemble from pentameric, hexameric, and pseudo-hexameric (trimeric) protein oligomers (*e*.*g*. Fig. 3A in Ref.^58^ and Refs.^18,19,179^). Experiments suggest that these oligomers are the basic assembly units, meaning that smaller protein complexes do not contribute significantly to the assembly process^19,180^. To focus on factors that control the composition of encapsulated cargo, we consider a minimal shell model developed in Perlmutter et al.^60^, with only pentamer and hexamer subunits that have interactions designed to drive assembly into a *T* =3 shells (containing 12 pentamers and 20 hexamers in a truncated icosahedron geometry). Our model builds on previous simulations of microcompartment assembly with no cargo^66^ or one cargo species^60,61,65^, as well as previous models for virus assembly^76,79,80,83,102,175–177,181–183^.

#### Shell-shell interactions

In our model, shell subunits interact through both repulsive and attractive forces. The repulsions consist of ‘Top’ (T) pseudoatoms that exist above the plane of the hexamer and ‘Bottom’ (B) pseudoatoms that exist below the plane. More specifically, we denote the top pseudoatoms as ‘TH’ and ‘TP’ respectively for hexamer and pentamer subunits, and similarly the bottom pseudoatoms as ‘BH’ and ‘BP’. All pairs of Top and Bottom pseudoatoms (excepting those on the same subunit) interact via a repulsive Lennard-Jones potential (Eq. B3). The Top-Top diameters and repulsion strengths set the shell spontaneous curvature and bending modulus (see section B).

Microcompartment protein shell assembly is primarily driven by interactions along the edges of the hexameric and pentameric subunits^58^. We represent these interactions by ‘Attractors’ on the perimeter of each shell subunit. Complementary Attractors on nearby subunits have short-range interactions modeled by a Morse potential (Eq. (B4) in section B). Attractors which are not complementary do not interact. The arrangement of Attractors on subunit edges is shown in Fig. 1, with pairs of complementary Attractors indicated by cyan double-headed arrows. The shell-shell binding affinity is proportional to the well-depth of the Morse potential between complementary Attractors, *ε*_SS_.

To focus on the role of interaction strengths in determining cargo encapsulation, we consider shell subunits that preferentially form only T=3 shells. To enforce this restriction, only three of the 6 edges of a hexamer have attractive interactions with pentamers, and there are no pentamer-pentamer attractive interactions (see Fig. 1).

#### Shell-cargo interactions

We model attractive interactions between hexamer subunits and the cargo particles by a Morse Potential between cargo particles and subunit ‘B’ pseudoatoms, with well-depth *ε*_SC_. These interactions represent the effect of ‘encapsulation peptides’ that target cargo to microcompartment interiors by mediating cargo-hexamer interactions^31,39,121,122,166^. To minimize the number of parameters, we keep *ε*_SC_ the same for both cargo species. We also add a layer of ‘Excluders’ in the plane of the ‘Top’ pseudoatoms, which represent shell-cargo excluded volume interactions. Since the shell-shell interaction geometries are already controlled by the Attractor, Top, and Bottom pseudoatoms, we do not consider Excluder-Excluder interactions.

#### Cargo-cargo interactions

In natural microcompartment systems, the interior cargo undergoes phase separation prior to or during shell assembly. The attractions between cargo particles that drive phase separation may be mediated by microcompartment scaffolding proteins (*e*.*g*.^120,122,126^) and/or direct pair interactions between cargo particles^36^. Similarly, in synthetic systems it is possible to engineer direct pair or scaffold-mediated attractions cargo molecules. In our model, we represent these scenarios in a minimal manner. We model both species of cargo particles with spherically symmetric excluded volume and direct pair attractions, implemented via an attractive Lennard-Jones (LJ) potential, with well-depth values *ε*_GG_, *ε*_RR_, and *ε*_RG_ for pairs of G-G, R-R, and R-G particles.

### B. Mapping simulation parameters to biological microcompartments

#### Relevant ranges of affinities

Although affinities among microcompartment components have not been measured, we estimate relevant ranges of our control parameters based on experimental observations of carboxysomes as follows.

##### Shell-shell affinities

Because empty carboxysome shells are not typically observed in cells (for wild-type carboxysomes) we have focused on shell-shell affinities that are below the threshold for empty shell assembly (*ε*_SS_ ≈ 4 *k*_B_*T*). Further, experimental observations suggest that *α*-carboxysomes assemble by one-step assembly pathways^57,120,161^, whereas *β* -carboxysomes^122,123^ and many other microcompartments^23,179^ assemble by two-step pathways. Therefore, except where mentioned otherwise, we consider two cases (*ε*_SS_*/k*_B_*T* = 2, 3.5), which are respectively below and above the transition between one-step and two-step assembly pathways (discussed in section II C).

##### Shell-cargo affinities

Since shell subunits robustly assemble around RuBisCO in both *α* and *β* carboxysomes, we have focused on shell-cargo affinities that are strong enough to observe this (*ε*_SC_ = 6 *k*_B_*T*, free energy *g*_SC_ ≈ − 14 *k*_B_*T*) except where mentioned otherwise.

##### Cargo-cargo affinities

For the case of *α*-carboxysomes, the RuBisCO does not undergo phase separation in the absence of shell proteins at physiological conditions^57,120,161^, but can phase separate without shell proteins under non-physiological conditions^126,128^. Thus, we infer that the effective cargo-cargo affinity for *α*-carboxysomes is near but below the threshold for phase separation, which corresponds to a cargo-cargo affinity of *ε*^vl^ ≈ 1.3 *k*_B_*T* at our simulated cargo concentration. In contrast, for *β* -carboxysomes the RuBisCO undergoes phase separation before shell assembly, suggesting a cargo-cargo affinity above *ε*^vl^, but below crystallization (*ε*^fs^). Our simulations span this range.

#### Length scales

Although the model is designed to be generic, we can *approximately* map it to carboxysomes by setting the cargo diameter (the unit length scale in the model) to that of the RuBisCO holoenzyme, implying *r* _*_ ≈ 13 nm. However, to enable tractable simulation of long assembly timescales, we have set the subunit/cargo size ratio to be larger than in carboxysomes — model subunits have a side length of *r*_*_ and are thus about three times larger than carboxysome hexamers (side-length ≈ 4 nm). See Ref. ^62^ for further discussion.

### C. Simulations and systems

We performed simulations using the Langevin dynamics algorithm in HOOMD (which uses GPUs to accelerate molecular dynamics simulations^184^), and periodic boundary conditions in a cubic box to represent a bulk system. The subunits were modeled as rigid bodies^185^. Each simulation was performed in the NVT ensemble, using the HOOMD fundamental units^186^, with the unit length scale *r*\* defined as the circumradius of the pentagonal subunit (the cargo diameter is also set to *r*_*_), and energies given in units of the thermal energy, *k*_B_*T*. The simulation timestep was 0.005 in dimensionless time units for all systems.

#### Systems

We simulated several systems as follows:

##### Shell assembly and cargo encapsulation

For dynamical simulations of shell assembly in the presence of cargo, each simulation contained enough subunits to form four complete microcompartments (48 pentamers and 80 hexamers), in a cubic simulation box with sidelength 40*r*\*. Each simulation also contained 854 cargo particles, with composition 50% ‘R’ and ‘G’ respectively. With *r*\* = 13 nm, this corresponds to concentrations of *c*_p_ = 0.6, *c*_h_ = 1, and *c*_c_ = 10 *μ*M for pentamers, hexamers, and cargo.

We performed most simulations for 2 × 10^6^ timesteps; simulations in the two-step assembly regime were performed for longer, 2 × 10^7^ timesteps, as they assemble more slowly.

##### Bulk simulations without shell subunits

We also performed large simulations of cargo without shell subunits, to determine the cargo phase behavior of the cargo as a function of the cargo-cargo affinities. Each of these simulations contained 46,938 cargo particles (50% ‘R’ and ‘G’ respectively) in a box with sidelength 160, to maintain the same cargo concentration as in the shell assembly simulations. We performed each simulation for 2 × 10^7^ timesteps.

##### Equilibrium cargo encapsulation

To identify effects of out-of-equilibrium dynamics of shell assembly on the properties of the encapsulated cargo, we performed additional simulations in which we simulated cargo dynamics in the presence of an assembled shell that was made permeable to cargo. In particular, these simulations consisted of a fully assembled immobile shell placed in the center of the box, with one missing pentamer to allow permeability to occur with the surrounding cargo. While the shell could not move, the *ε*_SC_ was still active. Each simulation contained 854 cargo particles with box sidelength 40*r*\* to maintain the same cargo concentration and composition as in the other systems. We performed each simulation for 2 × 10^6^ timesteps, which we determined was sufficient to achieve equilibration of cargo inside of the shell by monitoring the amount, composition, and mixing of the encapsulated cargo.

#### Initial conditions

Except for the equilibrium cargo encapsulation systems, all simulations were initialized with random positions for cargo particles and shell subunits (except that significant overlaps between pseudoatoms were forbidden). The equilibrium cargo encapsulation simulations were initialized with an assembled microcompartment shell whose subunits were fixed throughout the simulation, and the cargo particles were initialized with random locations with overlaps forbidden.

#### Statistical Significance

For most systems, we performed 10 independent trials at each parameter set. For most parameter sets at which we report statistical errors, we performed 20 independent trials. For parameter sets that resulted in low probability events (≤ 5%), we performed 60 trials to ensure low statistical error.

## Supporting information

Animation of assembly trajectory corresponding to Figure 3A

Animation of assembly trajectory corresponding to Figure 3B

Animation of assembly trajectory corresponding to Figure 3C

Animation of assembly trajectory corresponding to Figure 3D

Animation of assembly trajectory corresponding to Figure 3E

## ACKNOWLEDGMENTS

This work was supported by Award Number R01GM108021 from the National Institute Of General Medical Sciences and the Brandeis Center for Bioinspired Soft Materials, an NSF MRSEC, DMR-2011846. Computational resources were provided by NSF XSEDE computing resources (Maverick, Bridges, and Comet) and the Brandeis HPCC which is partially supported by DMR-2011846.

## DATA AVAILABILITY STATEMENT

The data that support the findings of this study are openly available in Open Science Framework at http://doi.org/10.17605/OSF.IO/SKZ7F

## Appendix A

Shell-shell and shell-cargo binding free energy values

We estimated the subunit-subunit binding free energy values as functions of the well-depth parameter *ε*_SS_ by measuring the dimerization equilibrium constant in simulations of subunits only capable of forming dimers. For both pentamer-hexamer and hexamer-hexamer dimers, we obtain binding free energies which are linear functions of the well-depth *ε*_SS_^60^. We interpret the y-intercept as the binding entropy. Setting *r*\* = 13 *nm* and using the conventional standard state concentration of 1 M (rather than 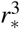 as in Ref. ^60^) gives

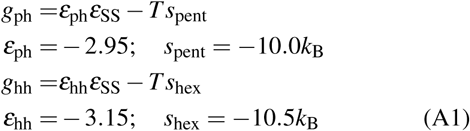

where *g*_ph_ and *g*_hh_ are the binding affinities for pentamer-hexamer and hexamer-hexamer pairs. The dissociation constants are then given by 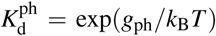 and 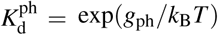. Table I shows the results for hexamer-hexamer dimerization, the values for pentamer-hexamer association are approximately the same. Similarly, we estimated the shell subunit adsorption free energy by performing simulations of subunits which cannot assemble (*ε*_SS_=0) in the presence of a cargo globule as^60^

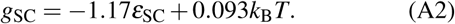

For the free energy of shell assembly, we consider a shell comprised of *n*_pent_=12 pentamers and *n*_hex_ hexamers, which have *n*_ph_ pentamer-hexamer contacts with binding energy *ε*_ph_ and *n*_hh_ hexamer-hexamer contacts with energy *ε*_hh_. For our *T* =3 model, *n*_hex_=20, *n*_ph_=60, and *n*_hh_=30. The assembly free energy is then given by

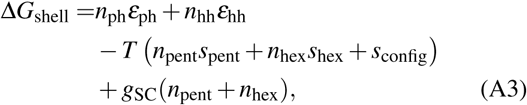

where *s*_config_ accounts for the configurational entropy associated with subunit and shell symmetries: 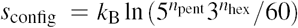, and *γ* and *A*_C_ are the surface tension and area of the cargo globule. Finally, we define an effective dissociation constant for the complex as 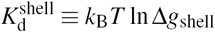 with Δ*g*_shell_ = Δ*G*_shell_*/*(*n*_pent_ + *n*_hex_) the free energy per subunit averaged over pentamers and hexamers (see Table I). However, this calculation is only approximate since it neglects changes in binding entropy between dimerization and the full complex. Further, to simplify the calculation and focus on the shell subunits, we have neglected cargo-cargo contributions to the complex stability. This provides a reasonable estimate for cargo-cargo affinities near liquid-vapor coexistence, but over/under estimates the shell stability for weaker/stronger cargo-cargo affinities. For a full analysis including cargo-cargo contributions, see Perlmutter et al.^60^.

## Appendix B

Model details

In our model, all potentials can be decomposed into pairwise interactions. Potentials involving shell subunits further decompose into pairwise interactions between their constituent building blocks – the Excluders, Attractors, ‘Top’, and ‘Bottom’ pseudoatoms. It is convenient to state the total energy of the system as the sum of three terms, involving shell-shell (*U*_SS_), cargo-cargo (*U*_CC_), and shell-cargo (*U*_SC_) interactions, each summed over all pairs of the appropriate type:

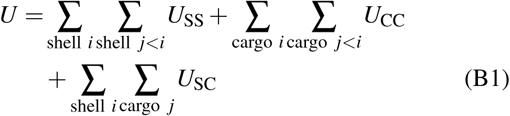

where Σ_shell *i*_ Σ_shell *j<i*_ is the sum over all distinct pairs of shell subunits in the system, Σ_shell *i*_ Σ_cargo *j*_ is the sum over all shell-cargo particle pairs, etc.

## Shell-shell interaction potentials

The shell-shell potential *U*_SS_ is the sum of the attractive interactions between complementary Attractors, and geometry guiding repulsive interactions between ‘Top’ - ‘Top’, ‘Bottom’ - ‘Bottom’, and ‘Top’ - ‘Bottom’ pairs. There are no interactions between members of the same rigid body. Thus, for notational clarity, we index rigid bodies and non-rigid pseudoatoms in Roman, while the pseudoatoms comprising a particular rigid body are indexed in Greek. For subunit *i* we denote its Attractor positions as {**a**_*iα*_} with the set comprising all Attractors *α*, its ‘Top’ position **t**_*i*_, and its ‘Bottom’ position **b**_*i*_.

The shell-shell interaction potential between two subunits *I* and *j* is then defined as:

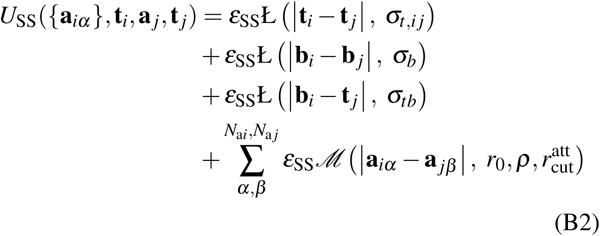

The function Ł is defined as the repulsive component of the Lennard-Jones potential shifted to zero at the interaction diameter:

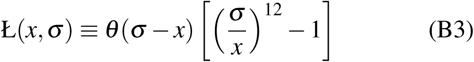

with *θ* (*x*) the Heaviside function. The function *ℳ* is a Morse potential:

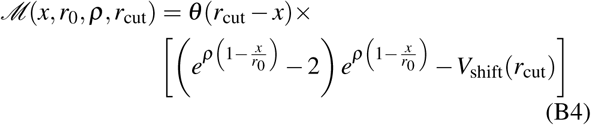

with *V*_shift_(*r*_cut_) the value of the (unshifted) potential at *r*_cut_.

The parameter *ε*_SS_ sets the strength of the shell-shell attraction at each Attractor site, *N*_a*i*_ is the number of Attractor pseudoatoms in subunit *i*, and *ε*_angle_ scales the repulsive interactions that enforce the geometry.

## Shell-shell interaction parameter values

### Attractors

The strength of attractive interactions is parameterized by the well-depth *ε*_SS_ for a pair of Attractors on hexamers as follows. Hexamer-Hexamer edge Attractor pairs (A2-A6, A3-A5, and A5-A6) have a well-depth of *ε*_SS_. Because vertex Attractors (A1, A4) have multiple partners in an assembled structure, whereas edge Attractors have only one, the well-depth for the vertex pairs (A1-A4 and A4-A4) is set to 0.5*ε*_SS_. Similarly, for pentamer-hexamer interactions, the well-depth for edge Attractor pairs (A2-A5, A3-A6) is *ε*_SS_, while the vertex interaction pairs (A1-A4 and A4-A4) have 0.5*ε*_SS_.

### Repulsive interactions

The ‘Top’ and ‘Bottom’ heights, or distance out of the Attractor plane, are set to *h* = 1*/*2*r*_b_, with *r*_b_ = 1 the distance between a vertex Attractor and the center of the pentagon. For all simulations shells have *T* =3 preferred curvature and,*σ*_*tb*_ = 1.8*r*_b_ is the diameter of the ‘Top’ - ‘Bottom’ interaction (this prevents subunits from binding in inverted configurations^76^), and *σ*_*b*_ = 1.5*r*_b_ is the diameter of the ‘Bottom’ - ‘Bottom’ interaction. In contrast to the latter parameters, *σ*_*t,i j*_ the effective diameter of the ‘Top’ - ‘Top’ interaction, depends on the species of subunits *i* and *j*; denoting a pentagonal or hexagonal subunit as ‘p’ or ‘h’ respectively, *σ*_t,pp_ = 2.1*r*_b_, *σ*_t,hh_ = 2.4*r*_b_, and *σ*_t,ph_ = 2.2*r*_b_. The parameter *r*_0_ is the minimum energy Attractor distance, set to 0.2*r*_b_, *ρ* = 4*r*_b_ determines the width of the attractive interaction, and 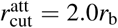 is the cutoff distance for the Attractor potential. Since the interactions just described are sufficient to describe assembly of the shell subunits, we included no Excluder-Excluder interactions.

## Cargo-cargo interactions

The interaction between cargo particles is given by a sum over all interacting pairs:

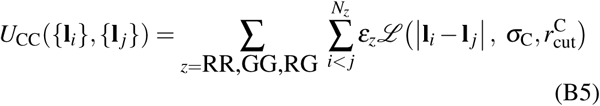

with *ℒ* the full Lennard-Jones interaction:

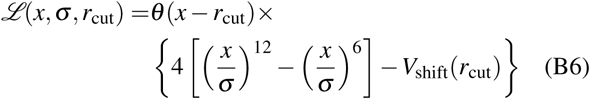

and *N*_RR_, *N*_GG_, and *N*_RG_ as the number of R-R, G-G, and R-G pairs in the system, and the cargo diameter *σ*_C_ = *r*_b_ and cutoff 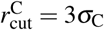 are the same for all cargo species.

## Shell-cargo interactions

The shell-cargo interaction is modeled by a short-range repulsion between cargo-Excluder and cargo-’Top’ pairs representing the excluded volume, plus an attractive interaction between pairs of cargo particles and shell subunit ‘Bottom’ pseudoatoms. For subunit *i* with Excluder positions **x**_*iα*_ and ‘Bottom’ psuedoatom **b**_*i*_, and cargo particle *j* with position **R** _*j*_, the potential is:

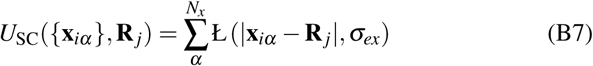

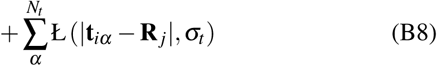

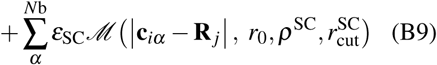

where *ε*_SC_ parameterizes the shell-cargo interaction strength, *N*_x_, *N*_t_, and *N*_b_ are the numbers of Excluders, ‘Top’, and ‘Bottom’ pseudoatoms on a shell subunit, *σ*_ex_ = 0.5*r*_b_ and *σ*_t_ = 0.5*r*_b_ are the effective diameters of the Excluder – cargo and ‘Top’ - cargo repulsions, 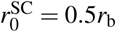 is the minimum energy Attractor distance, the width parameter is *ρ*^SC^ = 2.5*r*_b_, and the cutoff is set to 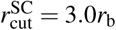.

